# Isolation and Genomic Characterization of *Desulfuromonas soudanensis* WTL, a Metal- and Electrode-Reducing Bacterium from Anoxic Deep Subsurface Brine

**DOI:** 10.1101/047530

**Authors:** Jonathan P. Badalamenti, Zarath M. Summers, Chi Ho Chan, Jeffrey A. Gralnick, Daniel R. Bond

## Abstract

Reaching a depth of 713 m below the surface, the Soudan Underground Iron Mine (Soudan, Minnesota, USA) transects a massive Archaean (2.7 Ga) banded iron formation, providing a remarkably accessible window into the terrestrial deep biosphere. Despite carbon limitation, metal-reducing microbial communities are present in potentially ancient anoxic brines continuously emanating from exploratory boreholes on Level 27. Using graphite electrodes deposited *in situ* as bait, we enriched and isolated a novel halophilic iron-reducing Deltaproteobacterium, *Desulfuromonas soudanensis* strain WTL, from an acetate-fed three-electrode bioreactor poised at +0.24 V (vs. standard hydrogen electrode). Cyclic voltammetry revealed that *D. soudanensis* releases electrons at redox potentials approximately 100 mV more positive than the model freshwater surface isolate *Geobacter sulfurreducens*, suggesting that its extracellular respiration is tuned for higher potential electron acceptors. *D. soudanensis* contains a 3,958,620-bp circular genome, assembled to completion using single-molecule real-time (SMRT) sequencing reads, which encodes a complete TCA cycle, 38 putative multiheme *c*-type cytochromes, one of which contains 69 heme-binding motifs, and a LuxI/LuxR quorum sensing cassette that produces an unidentified *N-*acyl homoserine lactone. Another cytochrome is predicted to lie within a putative prophage, suggesting that horizontal transfer of respiratory proteins plays a role in respiratory flexibility among metal reducers. Isolation of *D. soudanensis* underscores the utility of electrode-based approaches for enriching rare metal reducers from a wide range of habitats.

## INTRODUCTION

Microbial anaerobic respiration via metal reduction is a ubiquitous biogeochemical process in anoxic sediments and subsurface environments. Known metal reducers are found in several Bacterial and Archaeal phyla, and are often recognizable by their genomic arsenal of redox proteins, such as multiheme *c*-type cytochromes, that are implicated in transferring respiratory electrons across insulating biological membranes and cell walls to extracellular metal oxide particles. Many metal-reducing bacteria can also grow as biofilms using poised electrodes as terminal electron acceptors, a phenotype similar to the recently described phenomenon of syntrophic interspecies electron transfer (Rotaru et al., 2015). Because they can act as unlimited electron sinks, electrodes are powerful tools for enrichment of microorganisms capable of respiring extracellular acceptors. Mixed communities and pure cultures reducing electrodes have been obtained from habitats as diverse as wastewater sludge (Fu et al., 2013), freshwater and marine sediments (Bond et al., 2002; Holmes et al., 2004a; Miceli et al., 2012), soda lakes (Zavarzina et al., 2006), and deep subsurface fluids (Greene et al., 2009).

Along the southern edge of the Canadian shield, the Soudan Underground Mine in northern Minnesota (Figure 1A) transects the Archaean Animikie ocean basin (Ojakangas et al., 2001), with the main mineshaft providing access to 2.7 Gy-old banded iron deposits. On its lowest level (Level 27; 713 m depth), exploratory boreholes allow access to calcium- and metal-rich deep subsurface brines up to 3 times saltier than seawater, with low oxidation-reduction potentials, circumneutral pH levels, millimolar concentrations of reduced metals, and dissolved organic carbon concentrations below detection (Edwards et al., 2006; Toner et al., in prep.). Boreholes in the Soudan Mine, whose porewaters contain evidence of lithoautotrophic metabolism (Badalamenti et al., unpublished data), represent an opportunity to access a deep subsurface environment free of recent surface water contamination and photosynthetic inputs, in a region unperturbed by volcanic or tectonic activity.

**FIGURE 1.**
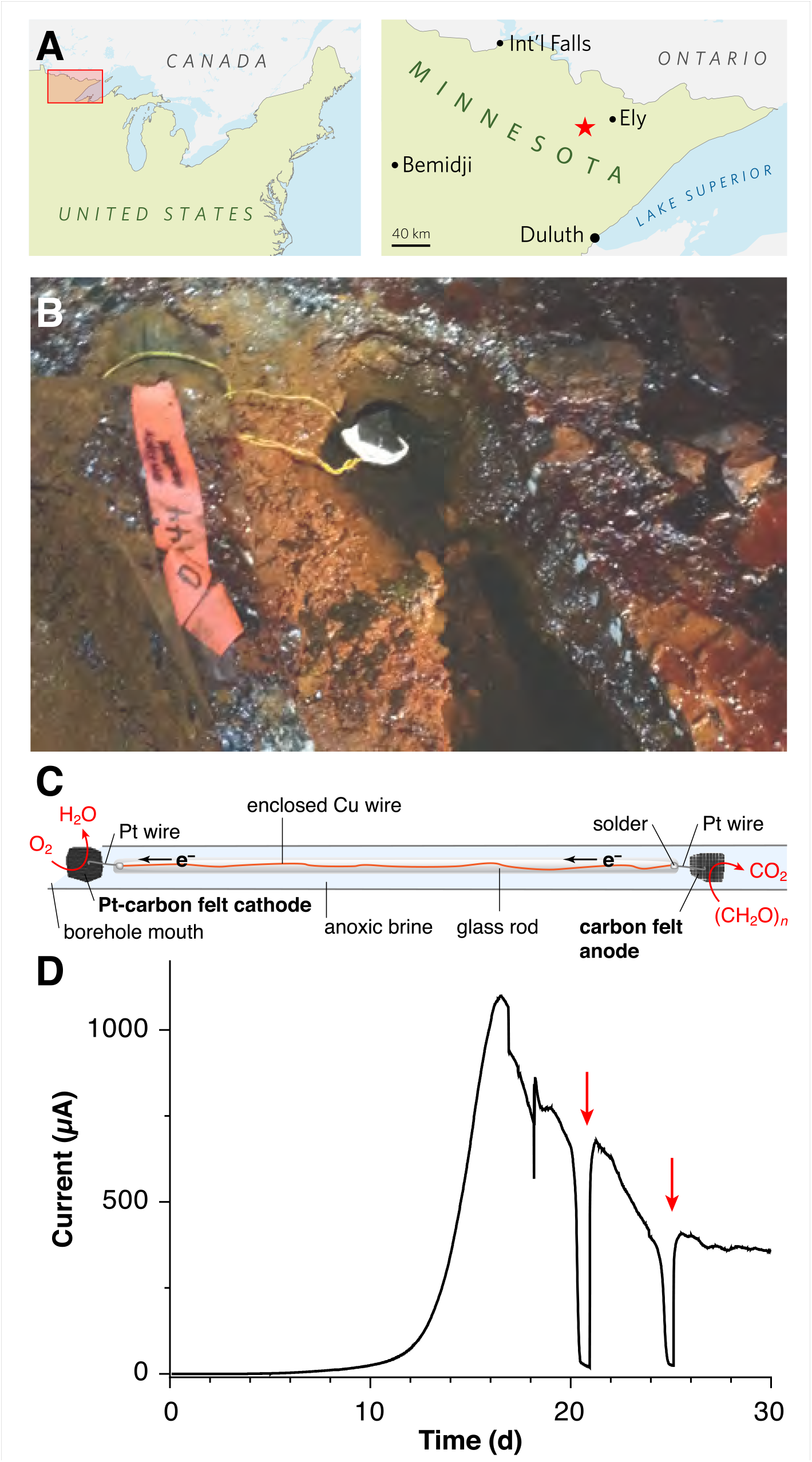
**Electrodes for enriching novel metal reducers from the terrestrial deep subsurface. A** - Map showing the location of the Soudan Underground Iron Mine within Minnesota’s Vermilion Range, approximately 30 km WSW of the town of Ely. **B** - Photograph of diamond drill hole (DDH) 944 located along the west tunnel of Level 27 (713 m depth below the surface). A white precipitate is visible coating the platinized carbon cathode. **C** - Schematic diagram of the electrode apparatus used for *in situ* enrichment. D - Chronoamperome-try of a potentiostatically controlled laboratory bioreactor inoculated with the anode from the *in situ* enrichment. Red arrows indicate addition of acetate following substrate depletion.

Here we describe isolation and phenotypic characterization of a metal-reducing subsurface isolate, *Desulfuromonas soudanensis* WTL, enriched by *in situ* electrodes placed in a Soudan Mine borehole. Measurements of Fe(III) reduction and electron transfer rates indicate that this isolate’s respiratory machinery is tuned for capturing energy from higher-potential electron acceptors than surface isolates. Its complete, circular chromosome is limited in cytochrome abundance compared to most described metal-reducing *Deltaproteobacteria* from surface sites or aquifers, and at least one of these cytochromes is embedded within a putative prophage, suggesting that virus-mediated horizontal gene transfer contributes to respiratory flexibility in the environment. The *D. soudanensis* genome encodes LuxI/LuxR quorum-sensing circuitry, and medium from *D. soudanensis* cultures activates an *N*-acyl-L-homoserine lactone (AHL)-sensing indicator strain (Zhu et al., 2003). These findings underscore the effectiveness of using microbial electrochemistry to obtain rare organisms from the deep terrestrial subsurface for reference genome sequencing and physiological characterization.

## MATERIALS AND METHODS

### Media and culture conditions

Soudan Mine medium (SM-1X) used for enrichment and isolation contained the following (per liter of deionized water): CaCl_2_ · 2H_2_O, 22.1 g; MgCl_2_ · 6H_2_O, 15.3 g; NaCl, 15.8 g; MgSO_4_ · 7H_2_O, 0.01 g; NH_4_Cl, 1.0 g; KH_2_PO_4_, 0.05 g; sodium acetate, 1.64 g; NaHCO_3_, 1.8 g; non-chelated trace minerals (Marsili et al., 2008), 10 ml; Wolfe’s vitamins, 10 ml. SM-0.5X medium, used for characterization and routine growth of the pure culture, contained half the concentration of chloride salts. When fumarate was the electron acceptor, fumaric acid was added to 40 mM final concentration and titrated to pH 6.0-6.1 with 50% (w/v) NaOH before adding other ingredients. When SM-0.5X medium was used for electrode-based growth of pure cultures, it contained 50 mM additional NaCl in lieu of fumarate. Fe(III)-reducing cultures contained either ferric citrate (55 mM), poorly crystalline iron oxide (~100 mM; Levar et al., 2014), or schwertmannite (~20 mM). Schwertmannite was prepared by addition of 5.2 ml 30% H_2_O_2_ to 1 l of 10 g/l FeSO_4_ · 7H_2_O and stirred overnight, and was collected by centrifugation followed by 3 rinses with deionized water. In all cases, Soudan Mine medium was prepared containing all ingredients except chloride salts and bicarbonate, brought to 0.7 times the final volume (i.e. 700 ml for 1 L), adjusted to pH 6.8 before adding NaHCO_3_, bubbled with oxygen-free 80:20 N_2_:CO_2_ and autoclaved in butyl rubber-stoppered tubes or serum bottles. A separate salt solution containing 73.5 g CaCl_2_ · 2H_2_O, 50.8 g MgCl_2_ · 6H_2_O, and 52.6 g NaCl per liter was bubbled with Ar and autoclaved separately. Upon cooling, this salt solution was aseptically and anaerobically added to basal SM medium to achieve the desired final volume.

### *In situ* electrode enrichment

A ~40-cm length of Cu wire was threaded into glass tubing, and soldered to Pt wire near the end of the hollow tube. The glass was fused shut at either end, leaving a short length of Pt wire exposed so that electrodes could be attached to each end and so that the whole device could be autoclaved and incubated in the mine without any copper wire exposure (Fig. 1C). One Pt lead was connected to a ~25-cm^2^ piece of graphite felt as the anode; the other was attached to an equal area of platinized carbon cloth (BASF Fuel Cell Co., Somerset, NJ) as the cathode. To remove impurities, electrodes were soaked sequentially in 70% EtOH, 1 N HCl, and 1 N NaOH, with diH_2_O rinses in between prior to autoclaving. The sterile rod containing the anode was slid gently down diamond drill hole (DDH) 944, located along the West Drift of Level 27 of the Soudan Underground Mine (Soudan, MN, USA; 47°49′24” N, 92°14′14” W) and held in place such that the cathode was exposed to the oxic opening of the borehole. After 2 months of incubation *in situ*, the anode was retrieved and small sections used to inoculate laboratory electrode enrichments as described below. Samples of DDH 944 borehole water were also brought to directly to the laboratory and used to inoculate reactors containing poised electrodes. However, out of 8 attempts (two separate sampling events), no laboratory reactors produced current above background levels after up to 60 days of incubation.

### Electrochemical methods

Anode sections from *in situ* mine enrichments were transferred to 100-ml glass cylindrical electrochemical cells (BASi, West Lafayette, IN) containing SM-1X electrode medium and 4 separate planar graphite electrodes (12 cm^2^ total surface area), a Pt counter electrode, and an Ag/AgCl reference electrode as previously described (Marsili et al., 2008). Electrodes were poised at +0.24 V vs. standard hydrogen electrode (SHE) with a potentiostat (Bio-Logic, Knoxville, TN) and bioreactors were incubated in a 20°C circulating water bath in the dark under constant magnetic stirring and flushing with humidified 80:20 N_2_:CO_2_ scrubbed free of O_2_. Cyclic voltammograms (CVs) were collected at 1 mV/s scan rate with the second of two scans reported. CVs were compared to those of *G. sulfurreducens* (Chan et al., 2015) grown at 30°C as previously described (Marsili et al., 2008).

### Isolation and cultivation

Enrichments were serially diluted in SM-1X medium containing 20 mM acetate and 100 mM Fe(III)-oxide, and the highest dilutions yielding Fe(II)-production at 24°C in the dark under an 80:20 N_2_:CO_2_ atmosphere were diluted further in 40 mM fumarate-containing medium. High dilutions (10^-6^) were streaked for isolation with fumarate as the electron acceptor in an anaerobic glovebag (Coy Laboratory Products, Grass Lake, MI) on vitamin-free SM-0.5X medium solidified with 0.9% (w/v) Bacto agar (Difco) amended with 0.5 mM cysteine, and incubated under a 75:20:5 N_2_:CO_2_:H_2_ atmosphere in an anaerobic jar (Almore International, Beaverton, OR) containing a basket of Pt catalyst recharge pellets (Microbiology International, Frederick, MD). After 3 weeks of incubation at 24 °C, isolated pink colonies were picked into 0.5 ml SM-0.5X medium containing 0.5 mM cysteine and subsequently transferred into 10-ml culture tubes. Possible contamination was assessed by fluorescence microscopy of DAPI-stained cells and by plating aerobically on either unbuffered SM agar or LB agar and anaerobically (75:20:5 N_2_:CO_2_:H_2_) on LB agar buffered with 1.8 g/l NaHCO_3_. SM-0.5X medium with 20 mM acetate and 40 mM fumarate was used for routine cultivation at 24°C. Growth with fumarate was determined by increase in OD_600_ and Fe(III) reduction was measured by the FerroZine assay. Axenic cultures were stored at −80°C in 10% (v/v) dimethyl sulfoxide, and a clonal population of revived *D. soudanensis* cells used to generate the genomic sequence was deposited in the German Collection of Microorganisms and Cell Cultures (DSMZ) as strain 101009. Because this enrichment survived unplanned oxygen exposures and temperature changes due to power outages, the isolate was dubbed strain WTL for “will to live.”

### Electron donor utilization

*D. soudanensis* cells grown with fumarate were inoculated (1:2 inoculum, as cells reached an OD600 of ~0.25) into magnetically stirred conical bioreactors containing a single graphite electrode poised at +0.24 V at 24°C as described previously (Marsili et al., 2008). To remove residual acetate, the potentiostat was paused and spent growth medium replaced twice with donor-free medium under a constant stream of 80:20 N_2_:CO_2_. Electrodes were then re-poised at +0.24 V for at least 30 min to obtain a steady background current in the absence of donors. The following electron donors were tested individually to 5 mM final concentration unless noted: lactate, ethanol, methanol, formate, glycerol, pyruvate, citrate, succinate, propionate, butyrate, glucose (2 mM), and benzoate (0.25 mM). If a substrate yielded an increase above background current, after 2 h medium was replaced twice again before testing another substrate; otherwise, up to 3 substrates were added cumulatively before 5 mM acetate was added to re-establish current production.

### Genome sequencing, assembly, and annotation

30 μg high MW genomic DNA (gDNA) was pooled from duplicate 10-ml stationary phase fumarate-grown cultures using a DNeasy Blood & Tissue kit (Qiagen). gDNA was quantified with Qubit (Life Technologies) and prepared for PacBio sequencing using standard 10-kb insert protocols. SMRT bell templates were size-selected at a 7-kbp cutoff using Blue Pippin electrophoresis (Sage Science). Long sequencing reads were collected from a total of 8 SMRT cells (P4-C2 chemistry, 120-min movies) on a PacBio RS II instrument (Mayo Clinic, Rochester, MN) yielding 1.7 Gbp of raw data (mean read length 4,663 bp; N_50_ 6,613 bp). Assembly was performed with HGAP v.3 (Chin et al., 2013) in SMRT Analysis v. 2.2 with a 10-kb minimum subread length to provide ~100x coverage. Unambiguous circularity of the resulting ~4-Mbp contig was confirmed via dot plot (Gepard v. 1.30) and self-complementary ends were manually trimmed. A crude initial annotation was generated with Prokka v. 1.10 (Seemann, 2014) to locate a single copy *of dnaA* upstream of several DnaA boxes predicted with Ori-Finder (Gao and Zhang, 2008). The manually reoriented contig was then polished using the entire PacBio dataset (~427x coverage) to QV > 50 before a final polishing step with 100x coverage of 250-bp paired-end Illumina reads using breseq v. 0.26 (Deatherage and Barrick, 2014) followed by Pilon v. 1.10 (Walker et al., 2014). The final 3,958,620-bp assembly was uploaded for automated annotation via the IMG/ER pipeline (https://img.jgi.doe.gov/cgi-bin/mer/main.cgi). Putative multiheme *c*-type cytochromes (≥ 3 Cxx(x)CH motifs) were identified with a Python script (https://github.com/bondlab/scripts), and Demerec gene abbreviations were assigned based on bi-directional best hits between *D. soudanensis* and *Geobacter sulfurreducens* PCA (GenBank accession no. NC_002939.5) as identified by GET_HOMOLOGUES (2015-05-29 release) (Contreras-Moreira and Vinuesa, 2013) at 30% identity and 90% query coverage. The manually curated IMG annotation was then submitted to GenBank (accession numbers provided below) to ensure consistency of genetic features across public databases.

### Phylogenomic classification and genomic analyses

FASTA nucleotide sequences for all publicly available *Desulfuromonas, Desulfuromusa, Geoalkalibacter, Geobacter, Geopsychrobater*, and *Pelobacter* genomes (26 total) were downloaded from NCBI or IMG/ER and analyzed with PhyloSift v. 1.0.1 (Darling et al., 2014). A concatenated alignment of 40 conserved single-copy marker genes was used to generate an unrooted maximum likelihood tree in FigTree v. 1.4.0 with *Desulfovibrio vulgaris* Hildenborough as the outgroup.

### Community analysis

Community gDNA harvested from unenriched Soudan brine (DDH 944), and a scraped section of electrode-enriched biofilms was isolated using a PowerWater Kit (Mo Bio, Carlsbad, CA). 16S rRNA genes were PCR amplified with phased V3-V4 primers as described previously (Bartram et al., 2011) and sequenced with 150-bp paired-end Illumina reads on a HiSeq 2000 (>22M reads/sample). Raw reads were quality trimmed, filtered, merged, aligned, clustered at 97% identity, and taxonomically assigned using mothur (Schloss et al., 2009) against the SILVA release 115 reference database. In addition, merged reads were mapped to the full-length 16S rRNA gene of *D. soudanensis* using bowtie2 v. 2.2.6 and mapped read counts were extracted from the resulting SAM files.

### Nucleotide accession numbers

Nucleotide sequence and annotation are available from GenBank (accession no. CP010802) and IMG/ER (GOLD analysis project Ga0069009). Raw reads and base modification data have been uploaded to the NCBI Sequence Read Archive under BioProject PRJNA272946.

## RESULTS

### Electrodes enrich rare taxa from subsurface brine

To enrich potential metal reducers within Soudan Mine boreholes, carbon cloth anodes were placed in the anoxic zone and connected to platinized cathodes in the oxic zone ~40 cm above, near the mouth of the borehole (Figure 1C). After 2 months of *in situ* incubation, 1 cm^2^ anode subsamples were transferred under anaerobic conditions to laboratory reactors containing acetate and poised graphite electrodes (+0.24 V, 20°C). Within 16 days, an exponential increase in anodic current, doubling every ~1 day was observed, reaching a maximum of 92 µA/cm^2^ (Figure 1D). Acetate addition after brief starvation periods immediately rescued current production (Figure 1D, red arrows), and transfer of scraped biofilm material from this initial enrichment to new reactors resulted in a similar growth pattern.

Amplicons from bacterial V3-V4 16S rRNA regions were generated to compare DNA recovered from the original borehole fluids with electrode-enriched communities. While *Deltaproteobacteria* sequences were among the least abundant lineages (<0.2%) in native borehole (DDH 944) brine, these sequences increased ~100-fold during laboratory electrode enrichments (20-22% relative abundance). In contrast, heterotrophic Marinobacter linages were equally abundant in both native and enriched samples (Bonis and Gralnick, 2015). With successive transfers, a Desulfuromonas sequence present at <0.02% relative abundance in the original anoxic fluid samples grew to comprise nearly 100% of the electrode Deltaproteobacteria. Other lineages linked to metal reduction, such as Acidobacteria (Mehta-Kolte and Bond, 2012) and the Peptococcaceae family of Firmicutes (Wrighton et al., 2011) were also present in brines, but they were not enriched. Inoculating borehole water directly into laboratory reactors did not produce any current-producing enrichments (n = 8).

### Pure cultures of *D. soudanensis* grow on electrodes and reduce Fe(III)

An isolate was obtained from electrode enrichments by first diluting in Fe(III)-oxide medium, then plating using fumarate as the electron acceptor, and was named *Desulfuromonas soudanensis* strain WTL. This strain demonstrated 96% full-length 16S rRNA gene sequence identity to *Desulfuromonas* sp. WB3 (enriched from a subsurface aquifer using As(V) as the electron acceptor; Osborne et al., 2015), *D. carbonis* (enriched from a coal bed gas well using Fe(III); An and Picardal, 2015), and *D. michiganensis* (enriched from river sediment using tetrachloroethene; Sung et al., 2003) (GenBank accession nos. KM452745.1, KJ776405.2, and NR_114607.1, respectively). Reads mapping with 100% identity to the 16S rRNA gene of the isolated *D. soudanensis* were present in the original DDH 944 brine DNA samples, albeit at extremely low relative abundance (data not shown). While original enrichments were conducted at 20°C and at salinities reflecting borehole conditions, *D. soudanensis* grew more rapidly at 24°C and at lower salt concentrations. These more favorable conditions were used for all subsequent characterization.

The maximum doubling time of *D. soudanensis* pure cultures growing on poised electrodes (+0.24 V vs. SHE) was 13.2 ± 0.05 h (*n* = 5). Based on current densities (58 ± 18 µA/cm^2^; *n* = 6), and attached protein levels (52 ± 11 µg/cm^2^; *n* = 5), the specific respiration rate of *D. soudanensis* was calculated to be 1.03 µA/µg protein (± 0.06; *n* = 5). This current:protein ratio, which was significantly slower than the 2-4 µA/µg ratios reported for *G. sulfurreducens* at various growth stages, agreed with the relatively slow doubling time of *D. soudanensis*. In addition, even with excess electron donor, biofilms reached a plateau at protein levels that suggested an inability to form multilayer biofilms on the electrode surface (10-20 µm thick biofilms typically correspond to attached protein of ~500 µg/cm^2^; (Marsili et al., 2008).

Cyclic voltammetry of *D. soudanensis* biofilms revealed a ~100-mV more positive shift in the potential triggering current flow from bacteria, as well as the midpoint, and saturation potentials of electron flux compared to *G. sulfurreducens* (Figure 2B). Single-turnover cyclic voltammetry resolved two major redox processes centered at −0.136 and −0.070 V vs. SHE, similar to other *Geobacter* biofilms where reversible oxidation/reduction midpoint potentials are slightly more negative than catalytic turnover voltammetry (Strycharz-Glaven and Tender, 2012; Marsili et al., 2010).

**FIGURE 2.**
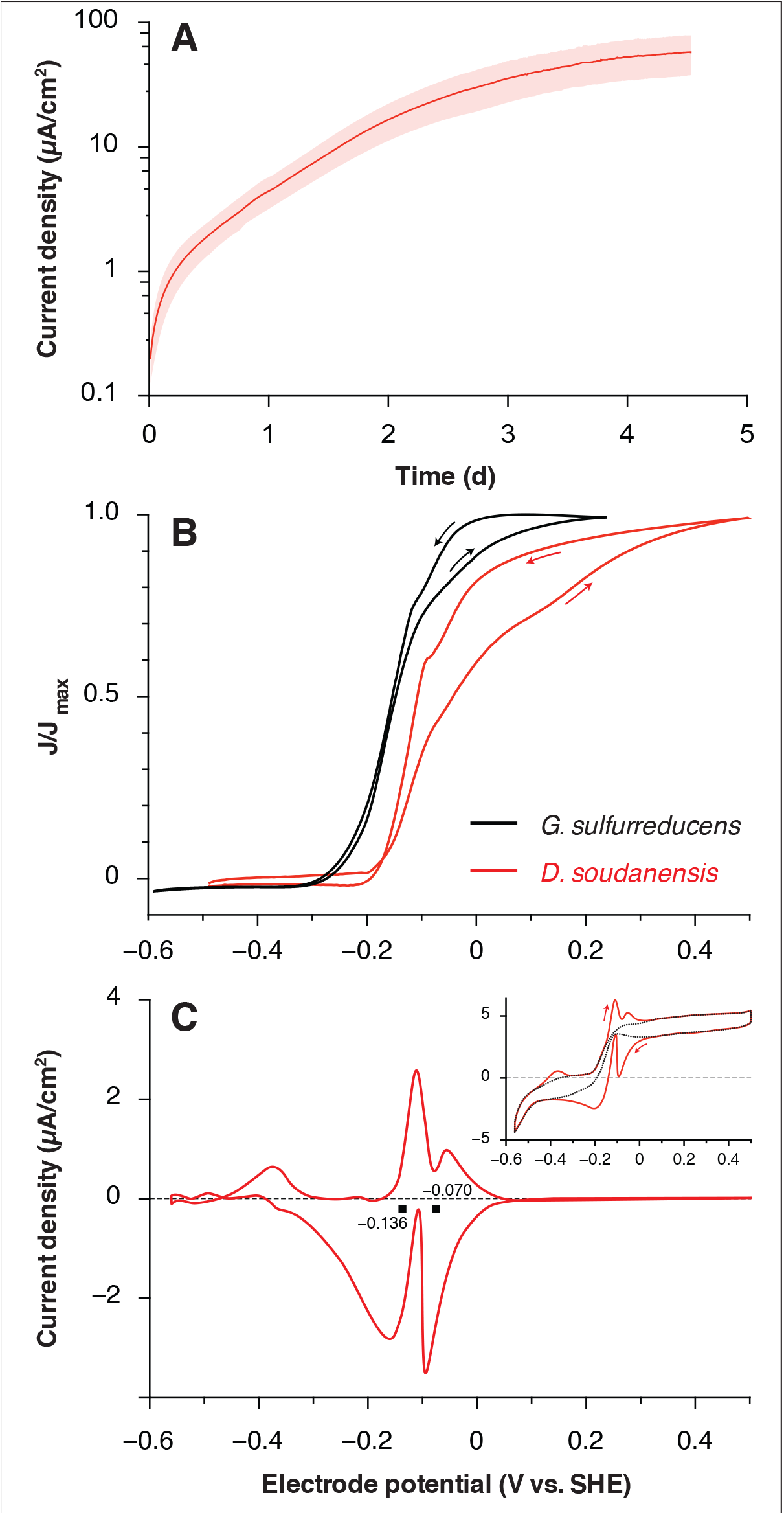
**Electrochemical characterization of pure cultures of *Desulfuromonas sou-danensis* WTL. A** - Chronoamperometry of potentiostatically controlled acetate-fed 10-ml bioreactors (+0.24 V vs. SHE) containing a single 3 cm^2^ graphite electrode. All electrode experiments were inoculated 1:2 with late log phase cultures pre-grown under electron acceptor limitation. The red trace and pink area show the mean and standard deviation, respectively, of 6 independent biological replicates. Estimates of doubling time were calculated against the linear portion of the curve from 0.7 to 2 d. **B** - Comparison of low scan rate (1 mV/s) cyclic voltammograms of *Geobacter sulfurreducens* (Chan et al., 2015; black trace) and *D. soudanensis* WTL (red trace). Current density on the *y*-axis is normalized against its maximum value for either organism. **C** – Baseline subtracted single-turnover cyclic voltam-mogram of an established *D. soudanensis* biofilm starved of acetate. Black squares denote the midpoint potentials of the two redox processes observed, and the inset shows the raw data (red trace) and polynomial function (black trace) used for baseline subtraction.

Electrochemical data showing a shift in the catalytic wave towards more positive redox potential suggested that *D. soudanensis* may be adapted to harvest energy from extracellular acceptors with higher redox potentials than those typically encountered by other bacteria previously characterized on electrodes. In experiments with insoluble Fe(III)-oxides, Fe(II) accumulation increased exponentially with a doubling time of 12.7 ± 1.1 h (*n* = 6) when freshly prepared schwertmannite, an acceptor with a predicted midpoint potential near +0.1 V SHE was provided (Thamdrup, 2000). Poorly crystalline Fe(III) oxide, predicted to have a redox potential near or below 0 V, supported a slower Fe(II) accumulation rate (15.7 ± 1.0 h, *n* = 3) (Figure 3B). With soluble acceptors, *D. soudanensis* showed poor Fe(III) reduction in ferric citrate medium, and growth slowed after accumulation of >3 mM Fe(II) (Figure 3C). With fumarate as the electron acceptor, cells grew exponentially, demonstrating a maximum doubling time of 13.3 ± 0.6 h (*n* = 3) when acetate was the electron donor, and fumarate fermentation was observed in donor-free controls (Figure 3D).

**FIGURE 3.**
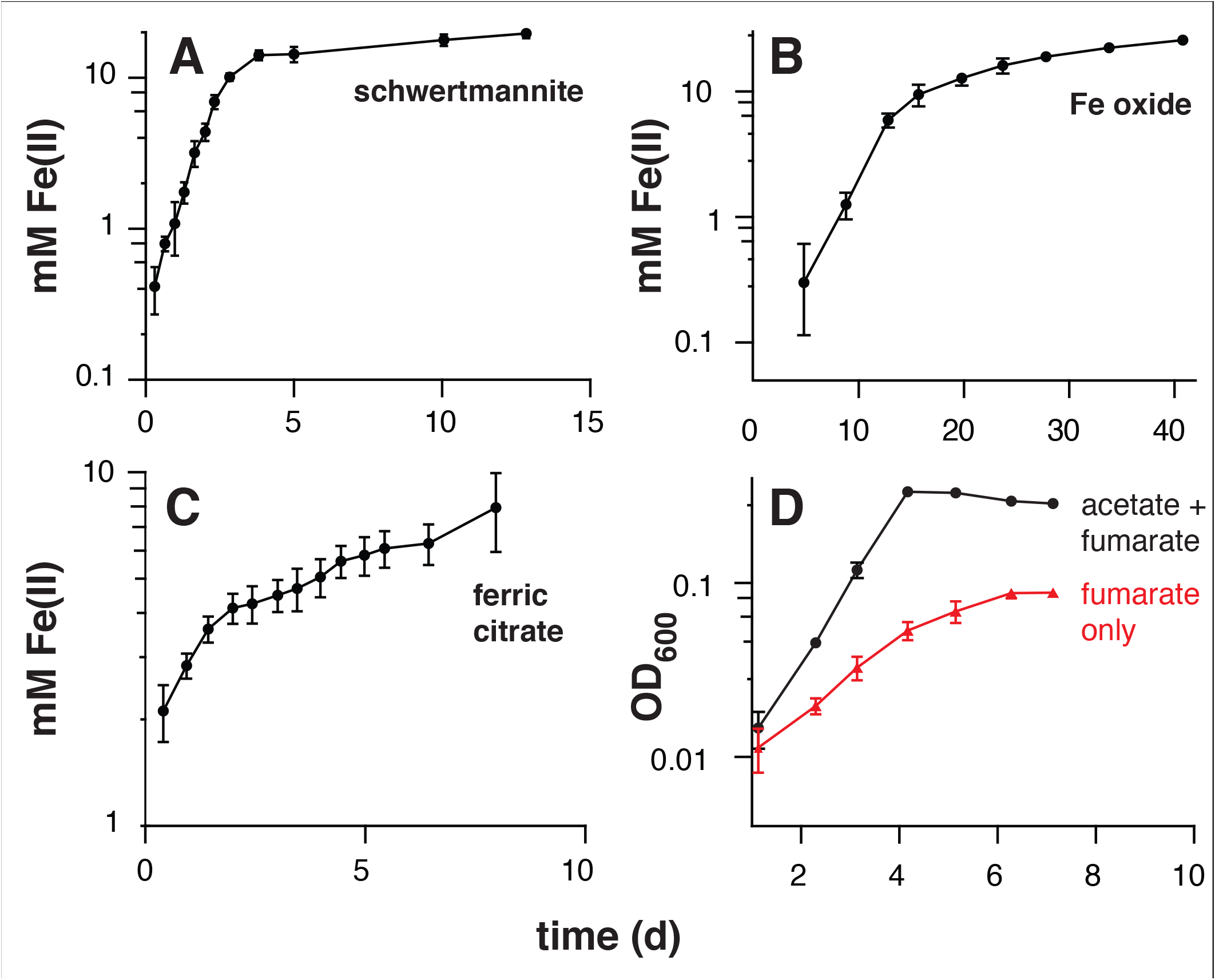
**Iron reduction and growth of pure cultures of *Desulfuromonas soudanensis* WTL on various electron acceptors with 20 mM acetate as the electron donor**. All plots are representative of three independent biological replicates. Accumulation of ferrous iron over time in incubations with either ~20 mM schwertmannite **(A)**, ~100 mM iron oxide **(B)**, or 55 mM ferric citrate **(C)**. In all cases Fe reduction slowed once Fe(II) concentrations reached 10 mM regardless of available Fe(III). **D** – Growth comparison of *D. soudanensis* cultures grown either with acetate plus 40 mM fumarate (black trace) or fumarate only (red trace).

### Genomic features

*D. soudanensis* contains a single 3,958-620-bp circular chromosome (Figure 4A) with similar G+C content (61.19%) to other *Desulfuromonas* spp., and represents the first complete *Desulfuromonas* genome. The genome encodes 3,419 protein-coding genes, 46 pseudogenes, 55 tRNAs, and 4 RNA polymerase sigma factors, and includes an exact tandem duplication of the 16S-5S-23S rRNA operon. Genes for assimilatory sulfate reduction, respiratory/dissimilatory nitrate reduction to ammonia, and N_2_ fixation are present, and *D. soudanensis* appears prototrophic for all amino acids, vitamins, and cofactors. Central metabolism in *D. soudanensis* includes a non-oxidative pentose phosphate pathway and Embden-Meyerhof-Parnas glycolysis/gluconeogenesis, linked by at least two putative pyruvate:ferredoxin oxidoreductases and one pyruvate dehydrogenase to a complete TCA cycle that includes the eukaryotic-like citrate synthase characteristic of other *Geobacter* and *Desulfuromonas* strains (Bond et al., 2005).

The *D. soudanensis* genome is dense with chemosensory and regulatory features, including 68 two-component histidine kinases, 97 transcriptional response regulators, 15 methyl-accepting chemotaxis proteins, 27 putative diguanylate cyclases and 16 predicted GEMM *cis*-regulatory cyclic di-GMP (or cyclic AMP-GMP) riboswitches. Among these, DSOUD_1479 bears highest identity (45%) to a recently reported diguanylate cyclase that produces c-AMP-GMP in *G. sulfurreducens* (Hallberg et al., 2016). Genes linked to growth in a metal-rich environment were also prevalent, such as an *hcgAB* gene cluster encoding mercury methylation (DSOUD_2468-2469; Parks et al., 2013), a Czc cobalt-zinc-cadmium exporter (DSOUD_2899-2901), CopA-family copper resistance system (DSOUD_1515-1516), and a cluster containing a glutaredoxin-dependent arsenate reductase, arsenite transporter, and an ArsR-family repressor (DSOUD_3330-3333).

A region encoding a LuxI-like acyl homoserine lactone synthase and a LuxR family transcriptional regulator is present in the vicinity of the encoded ImcH and CbcL homologs (Figure 4A), suggesting possible quorum-sensing abilities in this organism. Because production of AHL has never been described for any metal-reducing *Desulfuromonas* or *Geobacter* isolate, *D. soudanensis* was cultivated to ~0.5 OD_600_, cell-free supernatants were extracted with ethyl acetate, and dried extracts were resuspended in water. Extracts prepared from *D. soudanensis* cultures produced positive results for AHL-like compounds, using an indicator *Agrobacterium* strain (Zhu et al., 2003). Protein-normalized levels of beta-galactosidase activity resulting from addition of the *D. soudanensis* AHL were similar to positive control containing 10 nM *N*-3-oxohexanoyl-L-homoserine lactone, while an AHL-free control showed no activity (Figure 4C).

**FIGURE 4.**
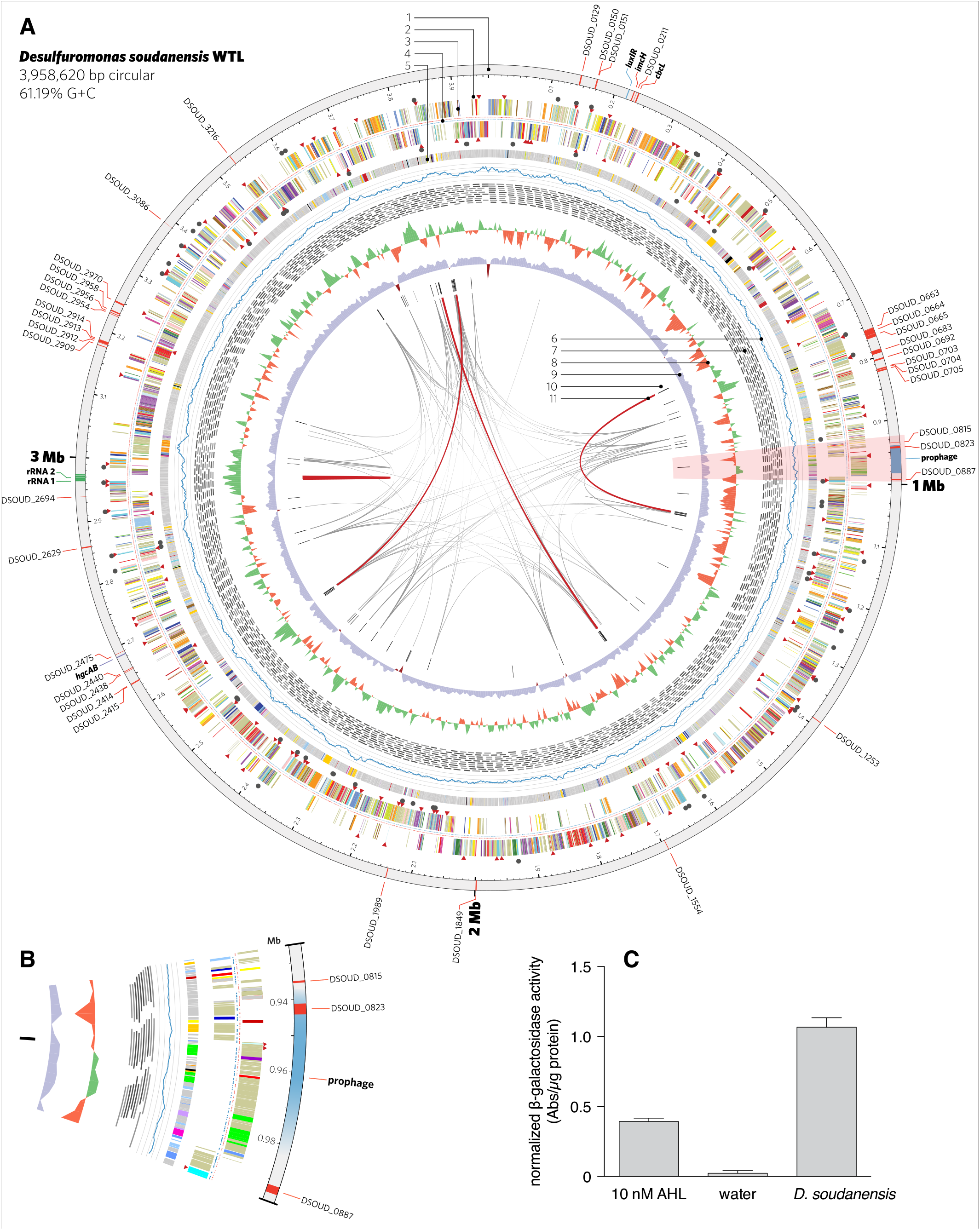
**Features of the complete genome of *Desulfuromonas soudanensis* WTL. A** -circular representation of the genome generated in Circos v. 0.64 (Krzywinski 2009). Rings are numbered moving from the outermost inward as follows: 1, location and locus tags of multiheme *c*-type cytochromes (red), rRNA operon duplication (green), and other features (blue); 2, putative sensor histidine kinases (red triangles) and response regulators (black circles); 3, protein coding sequences colored by COG category; 4, locations of methylated DNA bases identified by PacBio sequencing on the plus (red) and minus (blue) strands; 5, regions of putative horizontal gene transfer as predicted in IMG/ER, where grey is closest BLAST homology to other *Deltaproteobacteria*; 6, mapped read coverage (range 200-700x); 7, mapping positions of the longest reads in the PacBio dataset; 8, G+C skew in a 5-kbp windows; 9, G+C content (blue >50%; maroon <50%); 10, putative viral protein coding genes; 11, links showing repetitive sequence at 95% identity (grey, >500 bp; red, > 2 kbp). **B** - Zoomed region of the area highlighted in pink in Panel A showing the location of a putative prophage integrated into the *D. soudanensis* genome. **C** – Concentrated ethyl acetate extract of *D. soudanensis* supernatant triggers LacZ activity in an acyl-L-homoserine lactone indicator *Agrobacterium tumefaciens* strain (Zhu et al., 2003). LacZ activity in this strain is compared to a negative control (water) and to a positive control containing 10 nM authentic *N*-3-oxohexanoyl-L-homoserine lactone. Specific activity = ΔA_420_/ug protein.

### *D. soudanensis* carries a focused repertoire of multiheme cytochromes

*D. soudanensis* possesses 38 putative multiheme *c*-type cytochromes (3 or more Cxx(x)CH heme-binding motifs), a value about half of what is typically seen in *Geobacter* strains. Two of these proteins are homologs to inner membrane cytochromes ImcH and CbcL of *G. sulfurreducens* (DSOUD_0207 and DSOUD_0214; 39% and 71% BLAST identity, respectively), which have been implicated in transfer of electrons out of the quinone pool and into the periplasm (Levar et al., 2014; Zacharoff et al., 2016). Two inner membrane cytochromes were components of a putative NrfH/NrfA nitrite-to-ammonia respiration pathway. While most *Geobacter* species contain three to five triheme PpcA-like periplasmic cytochromes, only one (DSOUD_3086) was present in *D. soudanensis*. Only two outer membrane ‘conduit’ clusters consisting of multiheme cytochromes, lipoprotein cytochromes, and putative β-barrel proteins were identified (DSOUD_0702-0705, DSOUD_2909-2915), comprising 7 cytochromes. At least 4 cytochromes were predicted to have extracellular localization, and one of these (DSOUD_0664) contains 69 heme-binding motifs. One-third of the putative multiheme cytochromes identified in *D. soudanensis* had no obvious homologs in available databases, even when a recently obtained genome sequence was included (’*Ca*. D. biiwaabikowi’; Badalamenti et al., in prep).

Bioinformatic predictions using VirSorter (Roux et al., 2015) and phiSpy (Akhter et al., 2012) identified a region of the *D. soudanensis* genome bearing viral signatures such as a phage-like repressor and integrases, along with sheath, tail, and baseplate proteins. Within this predicted prophage genome, a large number of predicted horizontally transferred genes were present, and these were predicted to encode a triheme cytochrome, an 11-heme predicted lipoprotein cytochrome, and an MtrC family decaheme cytochrome (Figure 4B).

### Electrodes confirm genomic predictions for substrate utilization

When biofilms were starved and washed free of exogenous acetate, the real-time response to addition of various electron donors could be used as an indicator of the ability of *D. soudanensis* to metabolize that compound. Current increased immediately upon additions of lactate, ethanol, or pyruvate (Figure 5), consistent with genomic predictions for lactate dehydrogenase (DSOUD_0819), alcohol dehydrogenase (DSOUD_1067 and DSOUD_1075), and multiple mechanisms feeding pyruvate into the TCA cycle. H_2_ injected into the reactor phase also caused an increase in current (data not shown), in agreement with multiple [Ni-Fe] uptake hydrogenases encoded on the genome. In contrast, no electrochemical response was observed for methanol, glycerol, or glucose (Figure 5), agreeing with the lack of genes for activation, catabolism, and/or membrane transport of these substrates. Other substrates, including formate, citrate, succinate, propionate, butyrate, and benzoate were tested but failed to elicit a respiratory response within 30 min.

**FIGURE 5.**
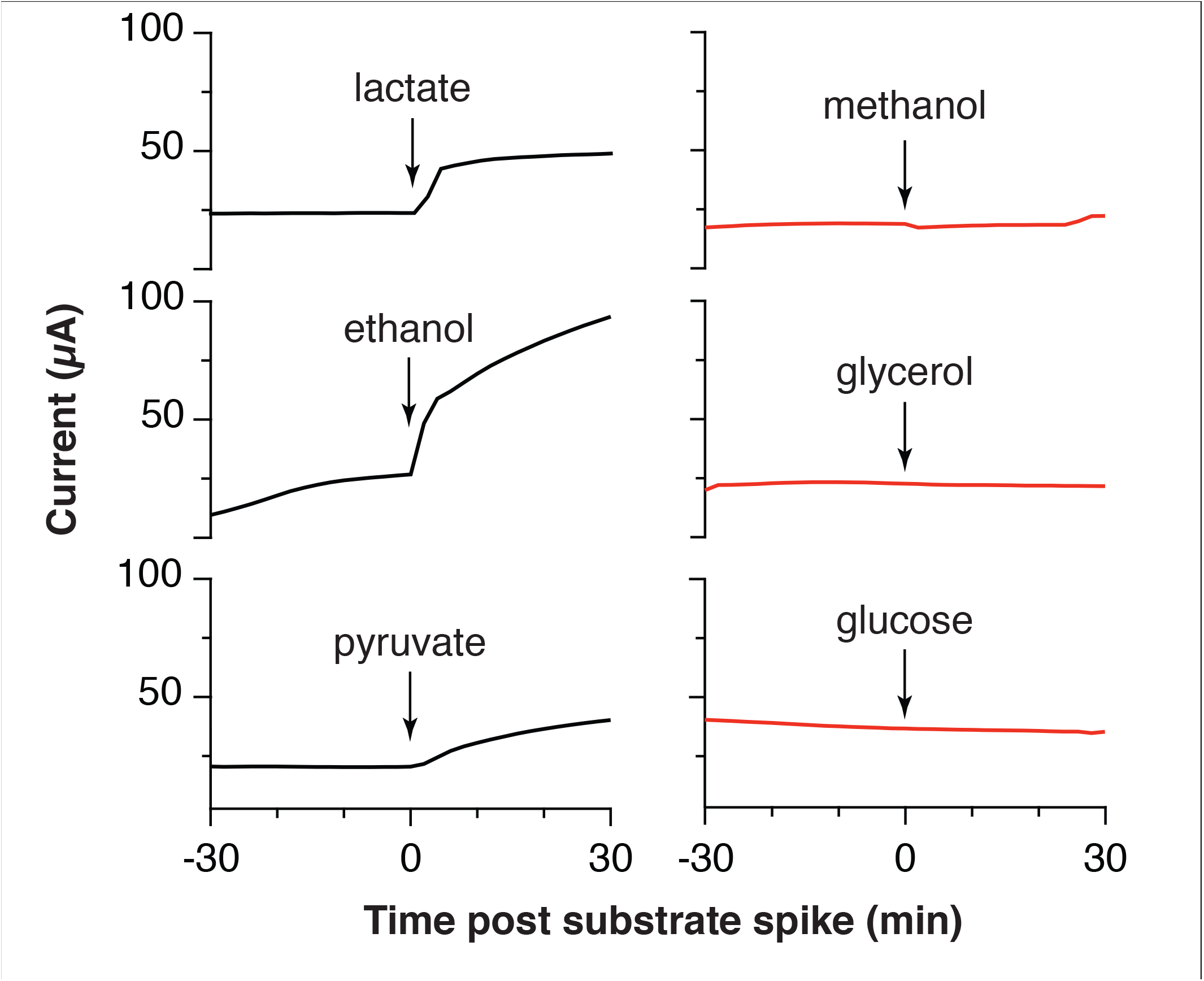
**Respiratory responses of starved, pre-established *Desulfuromonas soudanensis* WTL biofilms to addition of various electron donors**. All compounds were added to a final concentration of 5 mM except glucose (2 mM) into electrode bioreactors that were constantly poised at +0.24 V vs. SHE.

## DISCUSSION

By first enriching bacteria using anodes placed directly in anoxic Soudan Iron Mine borehole brine fluids, this study was able to isolate *D. soudanensis* WTL, a new metal- and electrode-respiring bacterium. In contrast, parallel attempts to directly inoculate laboratory reactors with borehole fluid samples consistently failed to obtain positive enrichments. As DNA surveys estimate *D. soudanensis* is present as less than 0.02% of organisms in borehole fluids, it is likely that samples from this ~10^3^ cell/ml environment contain, at best, a few *D. soudanensis* cells, even if they are concentrated via filtration. The rarity of this organism underscores the value of a narrow bottleneck such as that created by an electrode, which provides a consistent electron sink to support multiple doublings and increase cell numbers *in situ*.

Electrodes also enable real-time measurements of respiration rates in response to environmental or electrochemical perturbations, and this allowed screening a panel of utilizable electron donors (Figure 5) as well as surveying the redox potentials supporting extracellular respiration. The higher redox potential preferred by *D. soudanensis* (Figure 2B) is unique from characterized *Geobacter* strains, and provides evidence that the electron acceptor supporting growth of this organism in the environment is also different from what supports better-characterized freshwater isolates. *D. soudanensis* does have genes for the same inner membrane cytochromes (ImcH and CbcL) implicated in setting the redox potential preferences of *G. sulfurreducens*, raising the question of whether these cytochromes have altered redox potential windows, or if additional mechanisms exist (Levar et al., 2014; Zacharoff et al., 2016). Based on the recent observation that growth of thick multilayer biofilms on electrodes is correlated with the ability to form syntrophic associations with other bacteria (such as methanogens; Rotaru et al., 2015), the thin biofilms of *D. soudanensis* suggest a more isolated lifestyle.

Long read DNA sequencing has enabled cost-effective reconstruction of complete, reference-quality microbial genomes (Koren and Phillippy, 2015; Koren et al., 2013), even in this case where *D. soudanensis* possessed a tandem 16S-5S-23S rRNA operon duplication, as well as repetitive transposases, integrases, and recombinases (Figure 4A, internal links). The *D. soudanensis* genome represents the first complete genome from the *Desulfuromonas* genus, despite the type strain, *D. acetoxidans* DSM 684, having been first reported 40 years ago (Pfennig and Biebl, 1976). 278 proteins encoded in the *D. soudanensis* genome have no clear homologs (>30% threshold) in other metal-reducing *Deltaproteobacteria*, and among the 127 of these proteins with predicted functional annotation, a LuxI family *N*-acyl homoserine lactone synthase bearing only 24% BLAST identity to *G. uraniireducens* was identified. As *D. soudanensis* produces AHL-like compounds *in vivo* (Figure 4C), this finding suggests a previously undocumented role for quorum sensing in the broader context of Deltaproteobacteria, even in organic carbon-limited subsurface environments, where even small aggregates of *D. soudanensis* may need to respond to changes in flow rates or environmental conditions (Connell et al., 2010).

Though *D. soudanensis* only encodes roughly half as many *c*-type cytochromes as freshwater *Geobacter* spp., 11 of its 38 cytochromes are unique among known Deltaproteobacteria. Poor genomic conservation of outer surface cytochromes was first recognized ten years ago (Butler et al., 2010) and remains a pervasive pattern among metal-reducing bacteria in general. One hypothesis suggested by the *D. soudanensis* genome is that phage-mediated horizontal gene transfer contributes to cytochrome diversity. The closest homolog to the 11-heme cytochrome predicted to lie within a prophage (Figure 4B) is another cytochrome (33% amino acid identity) from *Geopsychrobacter electrodophilus*, an organism isolated from a marine sediment fuel cell in New Jersey, USA (Holmes et al., 2004a). When we analyzed all available metal-reducing Deltaproteobacterial genomes for viral signatures, *Gps. electrodophilus* was the only other genome that contained a cytochrome located within a putative prophage. Remarkably, both the cytochrome and prophage are different than those of *D. soudanensis*. Taken together, these findings suggest that viral transfer of cytochrome genes may be accelerating exchange of these cytochromes between metal-reducing bacteria. However, until phage particles can be recovered from these organisms, this remains a speculation.

The taxonomy of metal-reducing Deltaproteobacteria is still being resolved, particularly among isolates recovered from marine environments. While freshwater isolates are consistently named as *Geobacter* spp., marine isolates are often given genus designations that reflect isolation or enrichment conditions. Holmes et al. (Holmes et al., 2004b) proposed the family *Geobacteraceae* to include *Geobacter, Desulfuromonas, Desulfuromusa, Pelobacter*, and *Malonomonas*. Kuever et al. 2005); (corrig. 2006) described two family names with standing for classification in Bergey’s Manual, originally placing *Geobacter, Geoalkalibacter, Geothermobacter*, and *Geopsychrobacter* in the *Geobacteraceae* family, and assigning all other genera to the *Desulfuromonadaceae* family, based on designation of *D. acetoxidans* as the type strain (Pfennig and Biebl, 1976). Phylogenomic analysis supports an early evolutionary divergence between freshwater and marine strains (Figure 6), with 26 sequenced genomes separated into two distinct clades. Notably, this separation also tracks with per-genome cytochrome abundance, where freshwater strains contain much higher multiheme cytochrome content.

**FIGURE 6.**
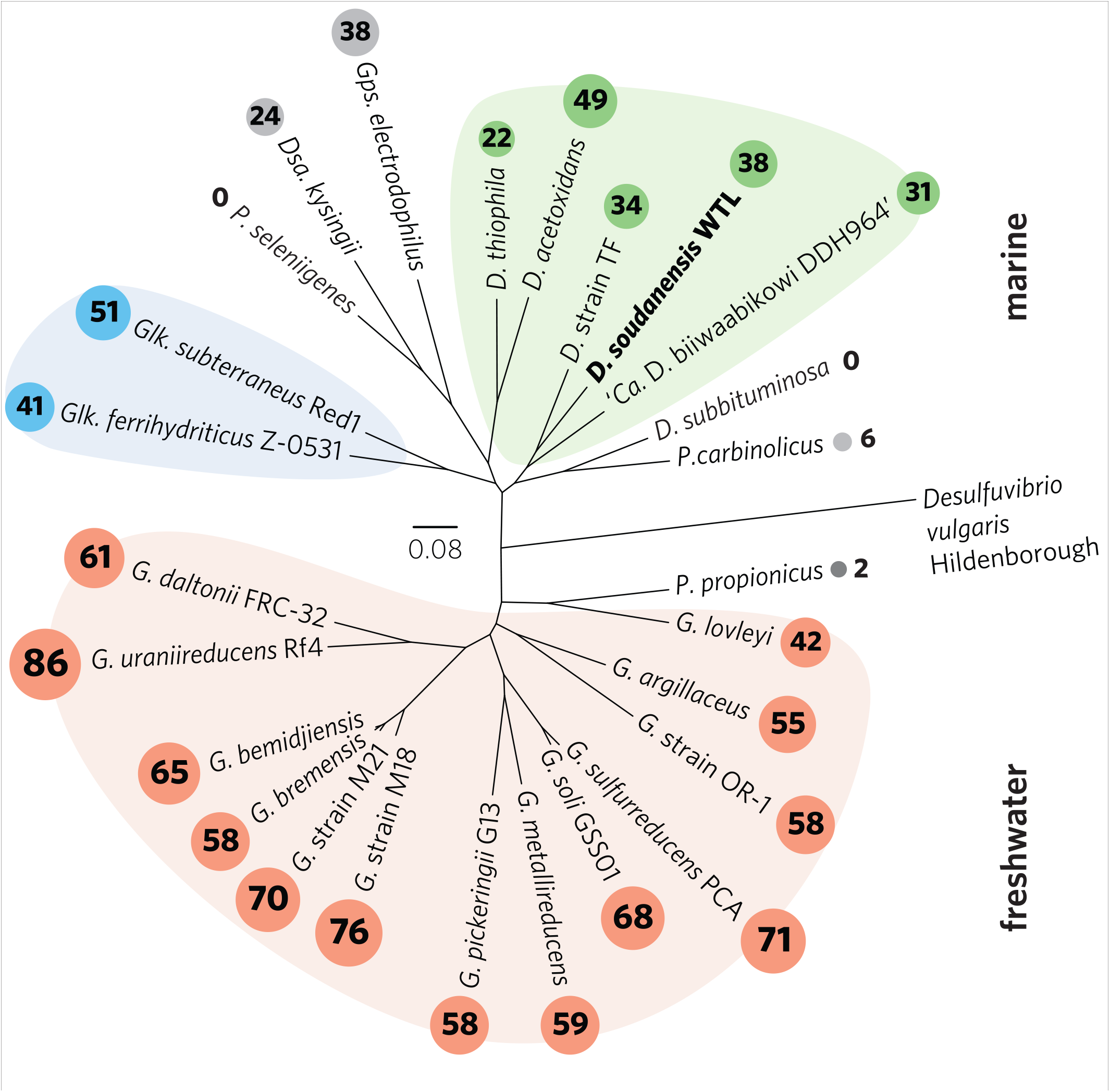
**Outgroup-rooted phylogenetic tree of *Geobacteraceae* and *Desulfuromonadaceae* family members with sequenced genomes**. The tree was constructed from an alignment of a concatenated set of 40 conserved single-copy marker genes in PhyloSift. Colored backgrounds are shown to group coherent genera: pink - *Geobacter*, blue - *Geoalkalibacter*, and green - *Desulfuromonas. D. soudanensis* WTL is shown in bold. Multiheme *c*-type cytochrome counts are shown in proportionally-sized circles. Genus abbreviations: *D*. -*Desulfuromonas, Dsa*. - *Desulfuromusa, G*. - *Geobacter, Glk*. - *Geoalkalibacter, Gps*. - *Geopsychrobacter, P*. - *Pelobacter*.

Based on whole-genome phylogeny, *D. soudanensis* falls into a clade with *Desulfuromonas* sp. TF (isolated using an electrode as the electron acceptor) and ‘*Ca*. Desulfuromonas biiwaabikowi DDH964’ a genome recovered from another Soudan electrode enrichment (NCBI BioProject Accession PRJNA316855; Badalamenti et al., in prep.). While this clade contains bacteria isolated from highly saline and subsurface sites, they are phylogenetically distinct from the two known halophilic *Geoalkalibacter* spp. (Greene et al., 2009; Zavarzina et al., 2006). The fact that each new metal-reducing isolate within this class shares a similar core anaerobic physiology, yet contains a highly variable collection of redox and sensory proteins suggests that many environmental niches defined by salinity, pH, and mineralogy are driving evolution and divergence within the *Geobacter-Desulfuromonas* cluster. Continued efforts to recover isolates and genomes from a wide diversity of habitats, using surfaces such as electrodes to target specific abilities, will aid in understanding the extent of this diversity.

## ACKNOWLEDGMENTS

We thank the Minnesota Department of Natural Resources and Jim Essig, Soudan Mine Underground State Park Manager, for providing logistical support during sampling trips. We thank Vuong Nguyen for performing the AHL LacZ activity assays. We also thank Karl Oles (Mayo Clinic Bioinformatics Core) for performing PacBio sequencing, and the Minnesota Supercomputing Institute for providing high-performance computing resources to support PacBio genome assembly and bioinformatics analyses.

